# Spatial domain analysis to estimate spatiotemporal pathological mechanisms in microenvironment with single-cell spatial omics data

**DOI:** 10.1101/2024.06.18.599475

**Authors:** Shunsuke A. Sakai, Ryosuke Nomura, Satoi Nagasawa, SungGi Chi, Ayako Suzuki, Yutaka Suzuki, Shumpei Ishikawa, Katsuya Tsuchihara, Shun-Ichiro Kageyama, Riu Yamashita

## Abstract

Single-cell spatial omics analysis requires consideration of biological functions and mechanisms in a microenvironment. However, microenvironment analysis using bioinformatic methods is limited by the need to detect histological morphology. In this study, we developed SpatialKNife (SKNY), an image-processing-based toolkit that detects spatial domains that potentially reflect histology and extends these domains to the microenvironment. The SKNY algorithm identified tumour spatial domains from spatial transcriptomic data of breast cancer, followed by clustering of these domains, trajectory estimation, and spatial extension to the tumour microenvironment (TME). The results of the trajectory estimation were consistent with the known mechanisms of cancer progression. We observed endothelial cell and macrophage infiltration into the TME at mid-stage progression. Our results suggest that analysis using the spatial domain as a unit reflects pathological mechanisms in the TME. This approach may be applicable to the biological estimation of diverse microenvironments.

## Introduction

Single-cell spatial omics platforms, such as Xenium, CosMx^1^, and PhenoCycler^2^, offer opportunities for the investigation of hundreds or thousands of genes in various organs and tissue types. The resolution of these methods is at the single-cell level, providing deep insight into the localisation of the expression of multiple genes in a particular microenvironment, which includes not only cancer cells but also immune cells and non-immune stromal cells. A key consideration in microenvironment analysis is the integration of gene expression and histological features to obtain a comprehensive understanding of biological functions and mechanisms. Classical methods that examine a microscope capture histological features through staining or fluorescence-based technologies, leading to the discovery of pathological mechanisms in the microenvironment^3^. However, in the current omics era, with the large number of specimens and gene panels, manual physical approaches are no longer sufficient.

To address the high throughput of omics data, several third-party tools such as Seurat and Scanpy have been developed to efficiently analyse expression data from thousands of gene panels and samples^4, 5, 6, 7, 8, 9, 10^. Methods inherited from single-cell RNA-seq have been implemented, including cell clustering^11, 12, 13, 14^, trajectory analysis^15, 16, 17, 18^, and ligand-receptor analysis^19, 20, 21^. These analytical methods use gene expression but do not consider molecular or cellular location. Hence, the integration of gene expression and location information is necessary for optimising spatial omics analysis of the microenvironment.

In response to this demand, several tools dedicated to spatial omics have been implemented, such as clustering analyses that integrate positional information with gene expression^22^ and ligand-receptor enrichment analysis at each spot in a space partitioned on a grid^23^. Although these methods are attractive for utilising spatial information, microenvironmental analysis is limited by the lack of direct histological information. Recently, the STARGATE algorithm^24^ was developed to detect spatial domains (i.e. regions with similar spatial expression patterns), and Sopa^25^ was constructed to extend ‘spatial domain’ analysis to single-cell spatial omics data. These methods can detect spatial domains that reflect and functionally resemble tumour, stromal, and vascular histologies.

Here, we extended the concept of the spatial domain to the microenvironment, which encompasses inside, peri-, and outside sections of the spatial domain, with the aim of estimating the functions and mechanisms of the microenvironment (Fig. 1a). We constructed an image processing-based toolkit, SpatialKNifeY (SKNY), to analyse the spatial domains from spatial omics data (Output 1-3) and extend it to the microenvironment (Output 4, 5) (Fig. 1b). Single-cell spatial transcriptomics data from Xenium^26^ was used to detect spatial domains of tumour for analysing the tumour microenvironment (TME) (Output 1: *Detection*) (Fig. 1c). Clustering of these spatial domains resulted in the formation of clusters consistent with the malignancy and subtypes (Output 2: *Clustering*), and the trajectory among spatial domains was estimated to represent the tumour progression process (Output 3: *Trajectory estimation*). The analysis extended from the spatial domain into TME and assessed the infiltration of endothelial cells into the tumour (Output 4: *Spatial stratification*). Furthermore, by integrating the trajectory and spatial stratification analysis, the dynamics in the tumour microenvironment were estimated, such as extracellular matrix degradation, angiogenesis, and macrophage infiltration (Output 5: *Spatiotemporal trajectory*). These results suggest that the SKNY can provide microenvironment analysis and may provide essential insights into their pathological functions. The SKNY algorithm is available under an open-source licence (https://github.com/shusakai/skny).

**Fig. 1.**
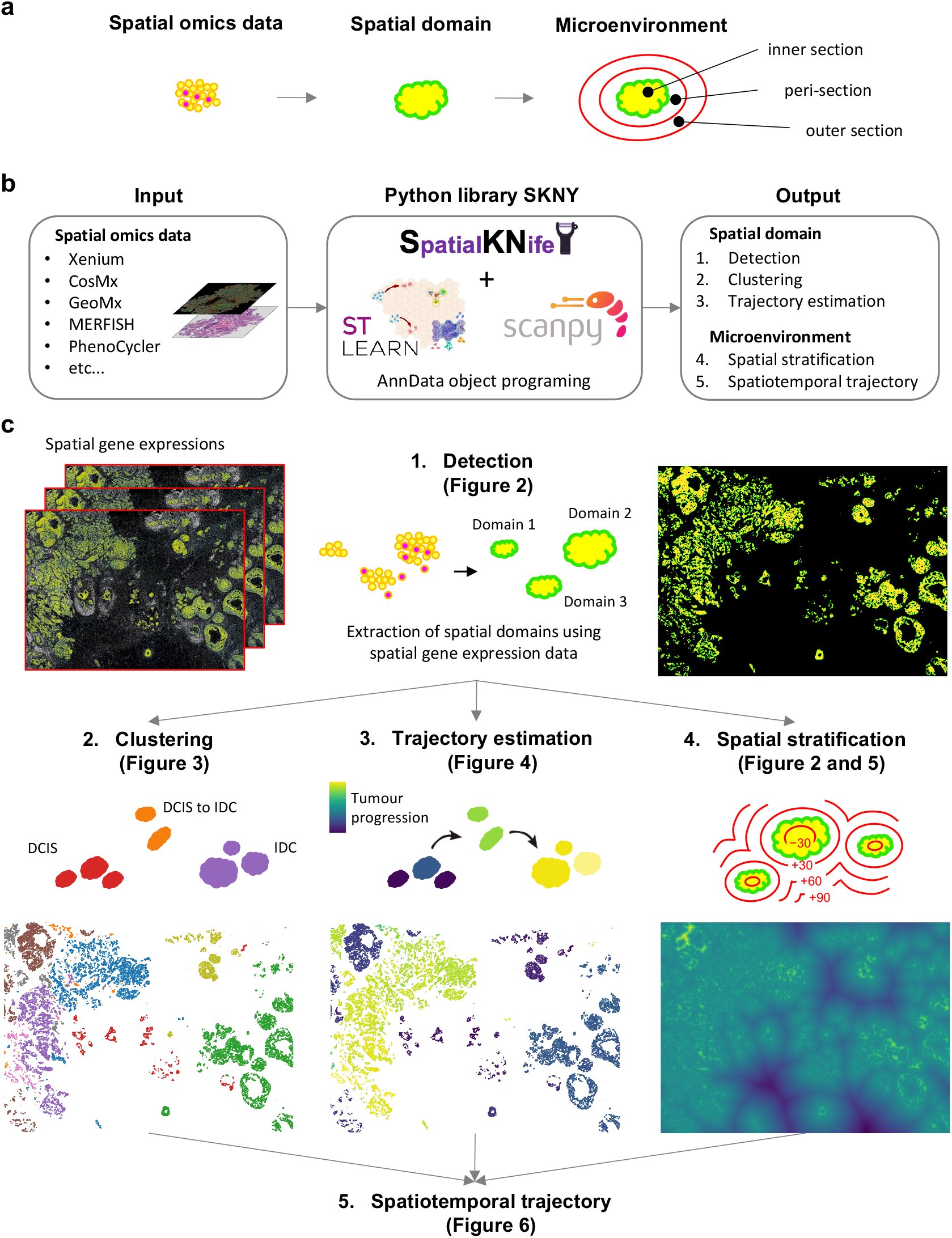
Landscape of SpatialKNifeY analysis. **(a)** The concept of the extension from spatial omics data and spatial domain to microenvironment. **(b)** The implementation of SpatialKNifeY (SKNY). A Python library of SKNY depends on stlearn^23^ and scanpy^9^ functions (see “Methods”) and AnnData object programing^10^. (c) The outputs from SKNY analysis. *Detection* (Output 1, see “Fig. 2”) delineates spatial domains based on a user’s positive and negative marker gene expressions. *Clustering* (Output 2, see “Fig. 3”) makes clusters of spatial domain units based on the mean expression of each gene. *Trajectory estimation* (Output 3, see “Fig. 4”) infers to the trajectory among spatial domains and pseudotime. *Spatial stratification* (Output 4, see “Fig. 5”) measures the distance from tumour boundary to each coordinate on the space and makes contour lines based on the distance. *Spatiotemporal trajectory* (Output 5, see “Fig. 6”) integrates pseudotime by *trajectory estimation* and contour lines by *spatial stratification* to estimate function and mechanism within the microenvironment.

## Results

### SKNY detects tumour spatial domains from Xenium data on breast cancer

To detect the spatial domain with the SKNY, we used Xenium data for breast cancer from a previous report^26^. A haematoxylin and eosin (HE)-stained image of the specimen from a previous report is shown (Fig. 2a). This specimen on a single slide contained various tumour tissues, including ductal carcinoma in situ (DCIS) and invasive ductal carcinoma (IDC). Using Xenium data, the SKNY algorithm was applied to detect tumour spatial domains (yellow) and extract their boundaries (green) based on the expression levels of the epithelial cell marker *CDH1* (Fig. 2b). Independently, the STARGATE algorithm^24^ was used to identify tumour spatial domains (Supplementary Fig. 1a, b, and c), resulting in high concordance with the results of SKNY (Jaccard similarity coefficient=0.85). The results suggest that the image-processing-based spatial domain extraction of the SKNY method is consistent with previous methods. The inward/outward areas from the extracted spatial domain boundaries were measured (Supplementary Fig. 2a), and the contour line was delineated at 30 µm intervals to spatially stratify the TME (Fig. 2c and Supplementary Fig. 2b). High-power field images, including single (Fig. 2c, left panel), triple (Fig. 2c, middle panel), and multiple spatial domains (Fig. 2c, right panel), showed visual concordance between the spatial domains and HE staining images for tumour detection.

**Fig. 2.**
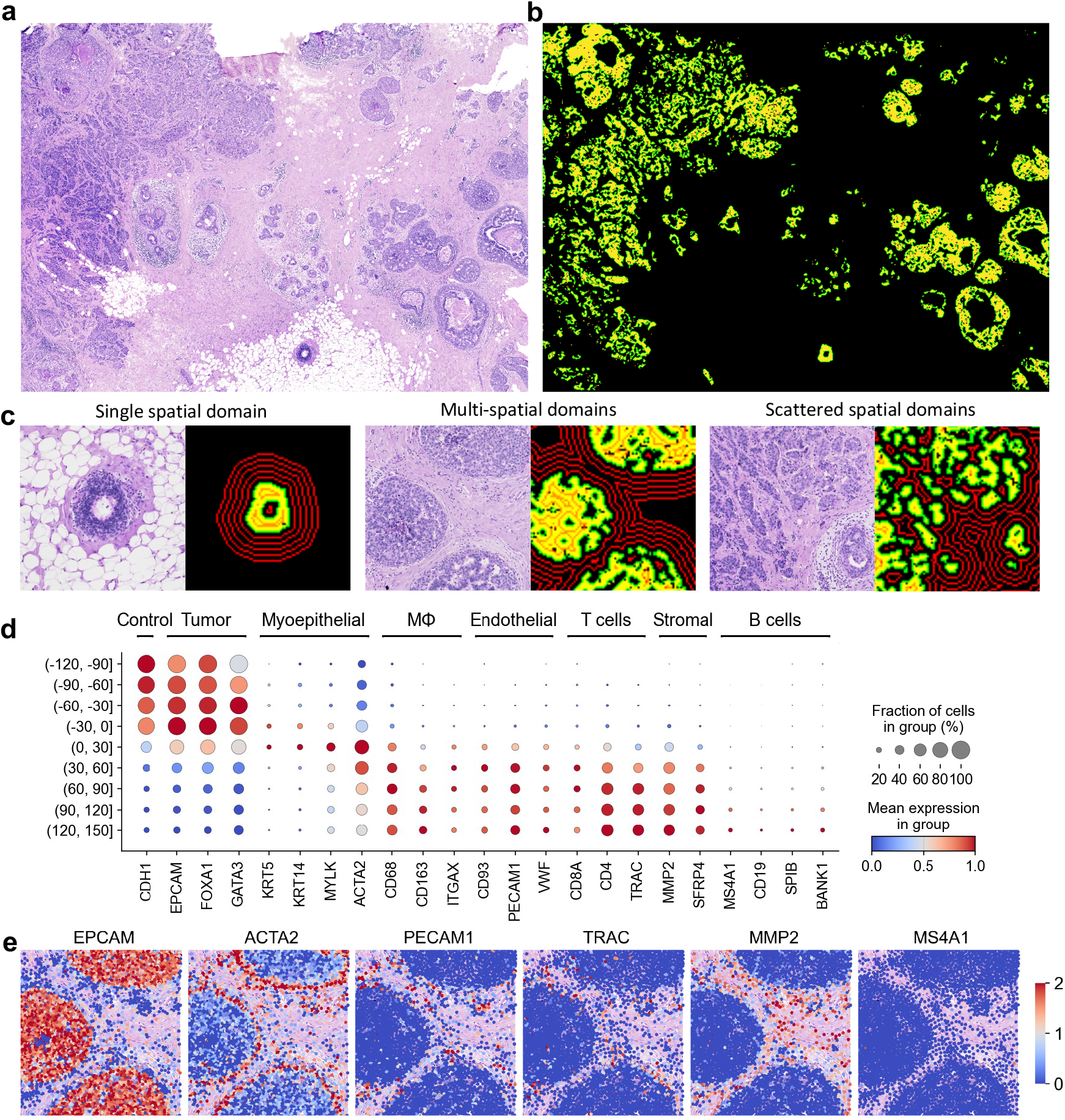
Detection of spatial domain with Xenium data accurately discriminates between the tumour and stromal region. **(a)** H&E staining image of breast cancer. **(b)** Detected spatial domains. The yellow and green colors indicate spatial domains and the boundary, respectively. **(c)** H&E staining images and spatial domain(s) from three ROIs. The red contour lines indicate distance from the surface of spatial domains at the interval of 30 μm. **(d)** Dotplot showing marker genes of each cell type. The color bar indicates the scaled mean count, and the size indicates the percentages of these gene expressions. **(e)** Spatial expression distribution of cell marker genes in the ROI. The color bar indicates the scaled mean count.

To confirm that these spatial domains are correctly partitioned between the tumour and stroma, the expression levels of several marker genes were examined in the stratified (−90, −60] to (+120, +150] sections in the total field. The results showed that cancer cell marker genes, such as *CDH1*, *EPCAM*, *FOXA1* and *GATA3* were enriched within the spatial domain (sections (-120, -90], (−90, −60], (−60, −30] and (−30, 0]) (Fig. 2d). The myoepithelial cell marker genes such as *KRT5*, *KRT14*, *MYLK* and *ACTA2* were enriched around the spatial domain boundary (the section of (0, +30]), and the macrophage, lymphocyte, endothelial cell, and stromal cell markers, such as *CD68*, *TRAC*, *PECAM1* and *MMP2*, respectively, were enriched on the outside (the sections of (+30, +60], (+60, +90], (+90, +120], and (+120, +150]). The spatial localisation of gene expression showed that *EPCAM* was overrepresented within the spatial domain, *ACTA2* at the boundary, and *PECAM1*, *TRAC*, and *MMP* outside the domain (Fig. 2e). These results suggest that the spatial domains stratified using the SKNY algorithm can be divided into tumours, peritumours, and stroma.

### SKNY clusters the spatial domains with multiple mixed cell types into subclusters using the UMAP algorithm

Next, to assess the diversity of cells within extracted spatial domains, we compared the α-diversity index (Chao1) based on the gene expression between cancer cells and spatial domains. The results indicated that the type of gene expression in the spatial domain was significantly more diverse than that in the cancer cells (*P*<0.001) (Supplementary Fig 3), suggesting that the spatial domains contain a variety of cells, not only cancer cells. Moreover, diversity variance was greater in spatial domains (standard deviation [SD]=62.1) than in cancer cells (SD=34.2). Hence, we hypothesised that the heterogeneity among spatial domains originated not only from cancer cells but also from diverse cells in the microenvironment. Here, we performed clustering of spatial domains to evaluate the heterogeneity among intra-spatial domain microenvironments. To obtain adequate gene expression data, 426 spatial domains larger than 1000 µm^2^ were selected. The gene expression data (313 genes) were dimensionally reduced by principal component analysis (PCA), resulting in nine clusters (0-8) based on their similarity in PCA space. Each spatial domain was placed in the two-dimensional space using UMAP (Fig. 3a) and the original space (Fig. 3b). To annotate these clusters with histology, we showed HE staining images based on the previous report (Fig. 3c). Combining this histology on HE staining with the clusters shown in Fig. 3b, we found that clusters 2, 3, 5, and 8 corresponded to non-invasive ductal carcinoma in situ (DCIS), whereas clusters 0, 1, 4, 6, and 7 corresponded to invasive ductal carcinoma in situ (IDC).

**Fig. 3.**
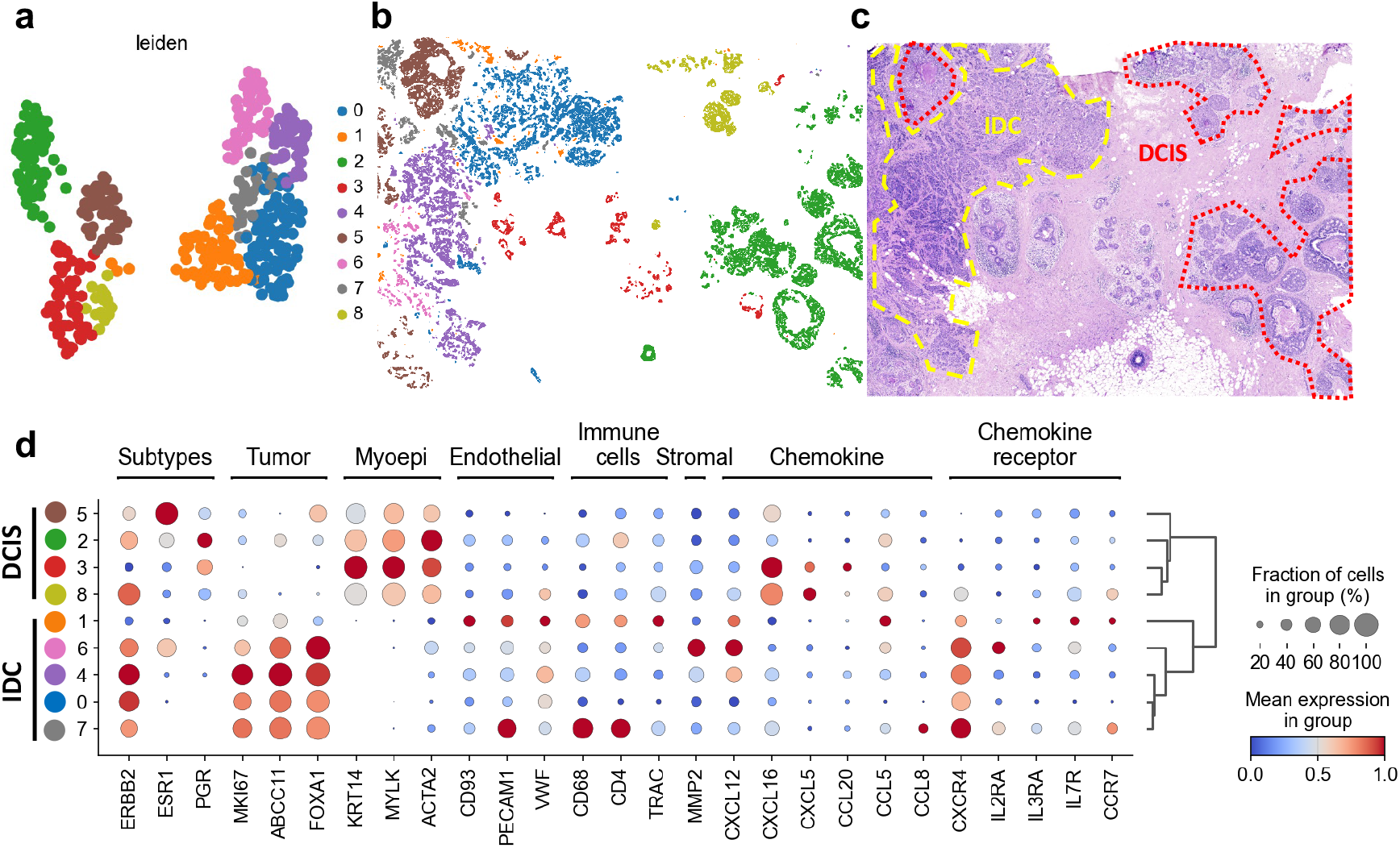
Clustering and annotation of spatial domain based on gene expressions. **(a)** Two-dimensional plot using UMAP loadings of gene expression of spatial domains. The colors indicate clusters. **(b)** Spatial distribution of each cluster. **(c)** H&E image with the histological annotations. **(d)** Dotplot showing markers of cell types and expression patterns of genes associated with tumour subtypes.

To provide detailed annotation of each spatial domain cluster, we examined the expression of several marker genes. In clusters 0, 1, 4, 6, and 7 (IDC clusters), *MKI67* and *ERBB2* were highly expressed. Conversely, in clusters 2, 3, 5, and 8 (DCIS clusters), the myoepithelial cell markers *ACTA2*, *MYLK*, and *KRT14* were highly expressed. These results suggest that gene expression in each spatial domain was consistent with the histological annotation (Fig. 3d). Interestingly, cluster 1 showed high expression of endothelial cell markers, including *PECAM1*, *VWF*, and *CD93*, as well as chemokines and chemokine receptor genes associated with cell migration, *CXCL12* and *CXCR4*. Furthermore, the expression of *MKI67*, *ABCC11,* and *FOXA1* was moderate in cluster 1 compared to that in other IDC clusters (Fig. 3d). Given the moderate expression of these cancer-associated genes and their midpoint in the UMAP space (Fig. 3a), Cluster 1 may represent a spatial domain at an intermediate stage in the transition from DCIS to IDC.

### SKNY estimates spatial domain trajectory, which reflects tumour progression

To estimate the spatial domain trajectory from DCIS to IDC, we used the partition-based graph abstraction (PAGA) algorithm^15^ to construct an adjacency graph representing the topology of expression patterns for each cluster (Fig. 4a). The adjacency graph is divided into clusters 2, 3, 5, and 8 (DCIS) and clusters 0, 4, 6, and 7 (IDC), where cluster 1 connects the DCIS and IDC clusters. Additionally, cluster 3, exhibiting the lowest tumour marker gene expression, as shown in Fig. 3d, was located at the lower end. This structure is consistent with the hypothesis that the spatial domain of DCIS clusters transitions to the IDC cluster via cluster 1. The pseudotime with cluster 3 as the root was determined and placed in the two-dimensional space of the PAGA algorithm and the original space (Fig. 4b left panel and Fig. 4c). We evaluated the correlation between the pseudotime and *MKI67* (*r*=0.52, *P*<0.001, Pearson coefficient)/*ACTA2* (*r*=−0.47, *P*<0.001). The pseudotime illustrated tumour progression (Fig. 4b middle and right panels).

**Fig. 4.**
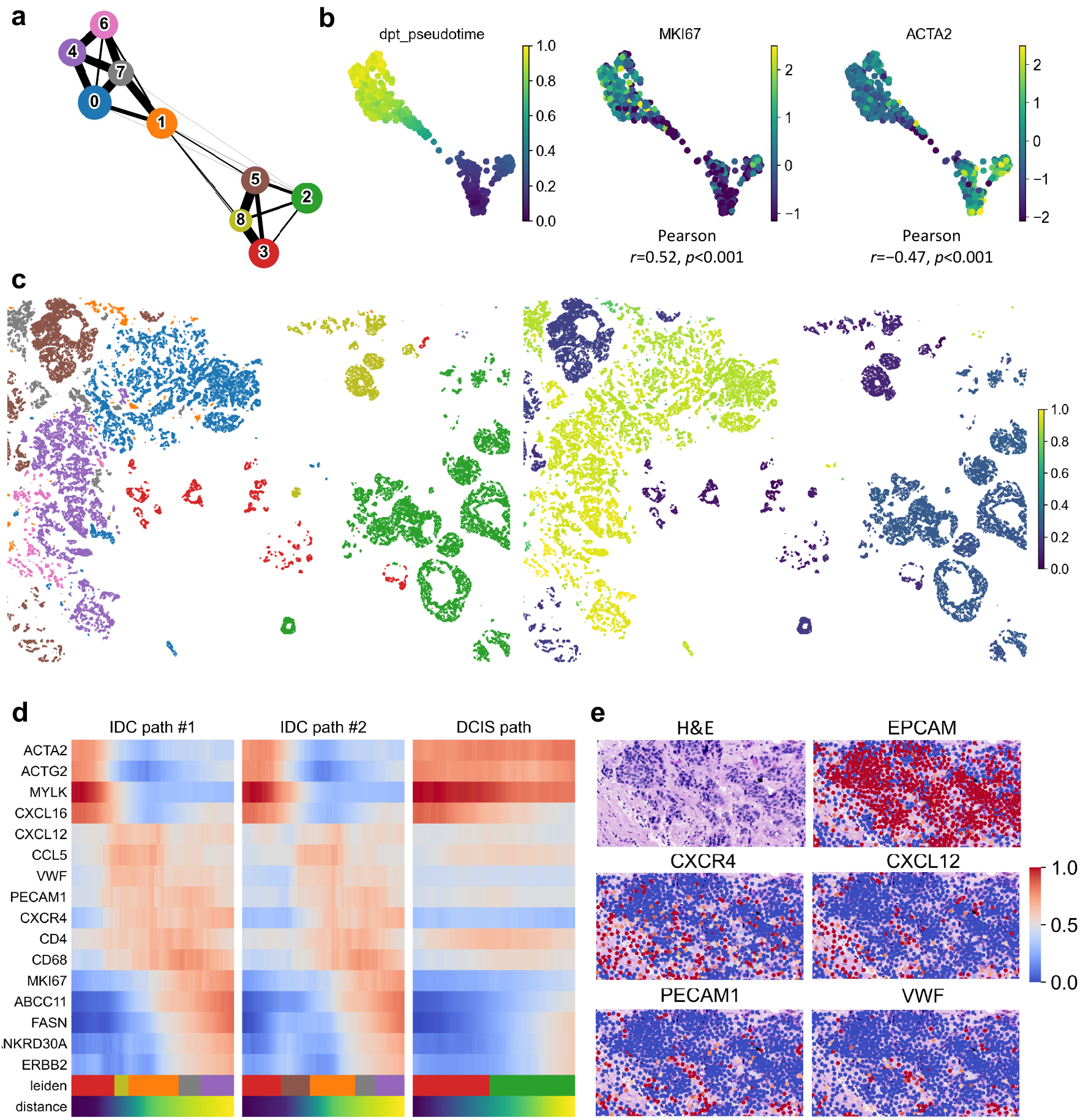
Estimating spatial domain trajectory reveals temporal gene expression gradient along with cancer progression. **(a)** PAGA graph constructed by the expression data of the spatial domains. **(b)** PAGA-initialized spatial domain embeddings with estimated pseudotimes, *MKI67*, and *ACTA2* expressions. Pearson’s correlation coefficients and the *P* values were used to evaluate the linear relationship between pseudotimes and scaled expression of *MKI67*/*ACTA2*. **(c)** Spatial distribution of clusters and pseudotimes. **(d)** Heatmap showing gene expression level along with pseudotimes on three progression paths. **(e)** Representative images of HE staining and gene expressions on the ROI. The color bar indicates the scaled mean count.

To identify characteristic gene expression at points on this pseudotime axis, we hypothesised three tumour progression paths (cluster 3→8→1→7→4: IDC path #1, 3→5→1→7→4: IDC path #2, and cluster 3→2: DCIS path) and evaluated trends in gene expression along these paths. In IDC paths #1 and #2, the expression of myoepithelial cell markers (*ACTG2* and *MYLK*) tended to decrease in the early stages of progression, whereas that of malignant markers (*ERBB2*) tended to increase in the later stages (Fig. 4d). In contrast, these myoepithelial cell and malignant marker fluctuations appeared to be moderate in the DCIS path. Moreover, in IDC paths #1 and #2, marker genes for endothelial cells (*VWF* and *PECAM1*), lymphocytes (*CD4*), macrophages (*CD68*), chemokines (*CXCL12* and *CCL5*), and chemokine receptors (*CXCR4*) were highly expressed at the intermediate stages of cancer progression. Similarly, in the DCIS path, *CD4*, *CD68*, and *CCL5* showed increased expression with progression. These findings suggest that endothelial cells and chemokine signalling are involved in the transition from DCIS to IDC. We also examined the spatial distribution of gene expression within the region of interest (ROI) corresponding to the transition phase from DCIS to IDC. The results showed a pattern in which *PECAM1*, *VWF*, *CXCR4*, and *CXCL12* appeared to infiltrate into regions of the tumour delineated by HE staining and *EPCAM* (Fig 4e). This also suggests that during the transition from DCIS to IDC, endothelial cells may infiltrate tumours and activate chemokine signals.

### SKNY quantifies the infiltrating of endothelial cells to spatial domains in the microenvironment

Next, we extended the spatial domains with their expression into their inner, peri-, and outer sections, namely, microenvironments, to quantitatively compare endothelial cell infiltration into tumours. We stratified the distance from the boundary of the spatial domain into 30 μm sections and extracted (−30, 0] (inner), (0, +30] (peri-), and (+30, +60] (outer) sections of each cluster (Fig 5a). Although no significant differences in expression levels were observed in the (+30, +60] section, significant differences among clusters were found in the (−30, 0] section for endothelial cell markers *PECAM1* and *VWF* (*P*=0.0053 and <0.001, Kruskal−Wallis test, respectively), with relatively high expression in cluster 1 (Fig 5b). To confirm the spatial expression patterns, ROIs selected from clusters 3, 8, 1, and 0 were extracted, and the distribution of cancer cell (*EPCAM* and *CDH1*) and endothelial cell (*VWF*, *PECAM*, *CD93*) markers was examined using Xenium Explorer. In clusters 3 and 8 (DCIS cluster), endothelial cell markers were localised outside the spatial domain, whereas in cluster 1 (DCIS-to-IDC cluster), these markers were localised in the tumour spatial domain (Fig. 5c). Moreover, cluster 0 (IDC cluster) appeared to remain in the gaps where the cancer cells had migrated (Fig. 5c, right panel). These results demonstrate that the analysis, expanded from the spatial domain to the microenvironment, could reflect the infiltration of endothelial cells into the tumour.

**Fig. 5.**
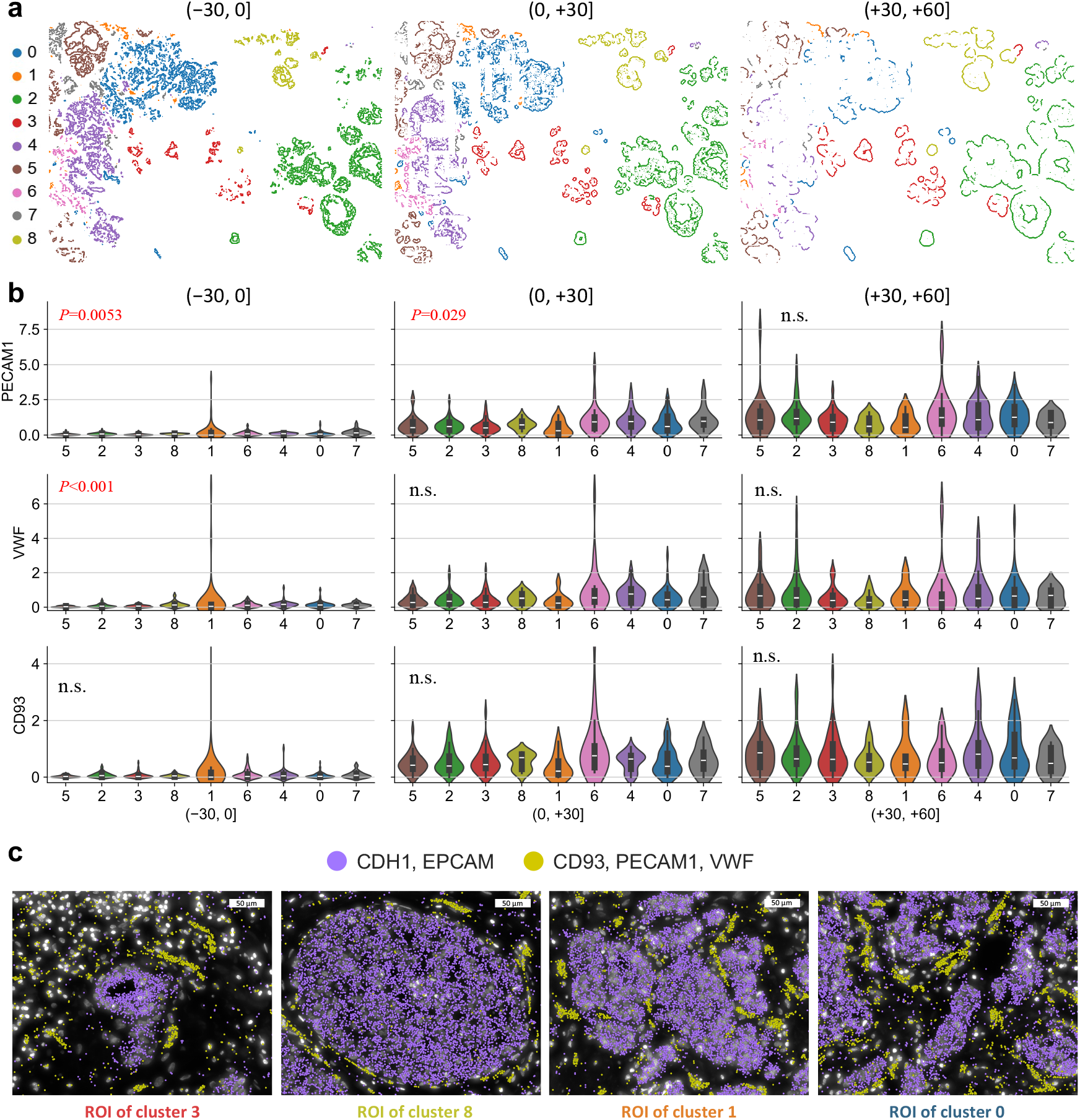
Spatial stratification of each spatial domain cluster elucidating endothelial cell invasion into the tumour. **(a)** Spatial distributions of stratified spatial domain clusters into (−30, 0], (0, +30], and (+30, +60] sections. **(b)** Violin plots showing the endothelial cell marker gene expressions (*PECAM1*, *VWF*, and *CD93*) for each cluster in the (−30, 0], (0, +30], and (+30, +60] sections. The x-axes indicate cluster numbers, and the y-axes indicate scaled gene expression levels. The annotated values are the *P* values of the significance test. **(c)** Representative images of DAPI with epithelial cell markers (*CDH1* and *EPCAM*) and endothelial cell markers (*CD93*, *PECAM1*, and *VWF*) expressions for four ROIs.

### *SKNY performs* spatiotemporal *trajectory analysis and estimates the mechanism of tumour progression*

Next, we analysed the spatiotemporal dynamics of gene expression by integrating the spatial axis of the microenvironment with the temporal axis estimated from the trajectory of tumour progression. We examined the changes over the pseudotime (IDC path #1) in the expression of endothelial cells (*PECAM1*), macrophages (*CD68*), matrix metalloproteinases (*MMP2*), chemokine receptors (*CXCR4*), and chemokines (*CXCL12*) in each TME section at (−30, 0] (inner), (0, +30) (peri-), and (+30, +60] (outer), respectively. In the inner section, *PECAM1* (*P*=0.023), *CD68* (*P*<0.001), and *CXCR4* (*P*<0.001) showed an increase during the transition period from DCIS to IDC (clusters 8, 1, and 7), whereas in the peri-section, *PECAM1* (*P*=0.035) and *CXCR4* (*P*<0.0024) also showed an increase during that period (Kruskal-Wallis test, Bonferroni-corrected *P* values) (Fig. 6a). In contrast, in the peri- and outer sections, *MMP2* (*P*<0.001 and *P*=0.020, respectively) showed an increase in these peaks in early DCIS (cluster 3) and late IDC (cluster 4).

**Fig. 6.**
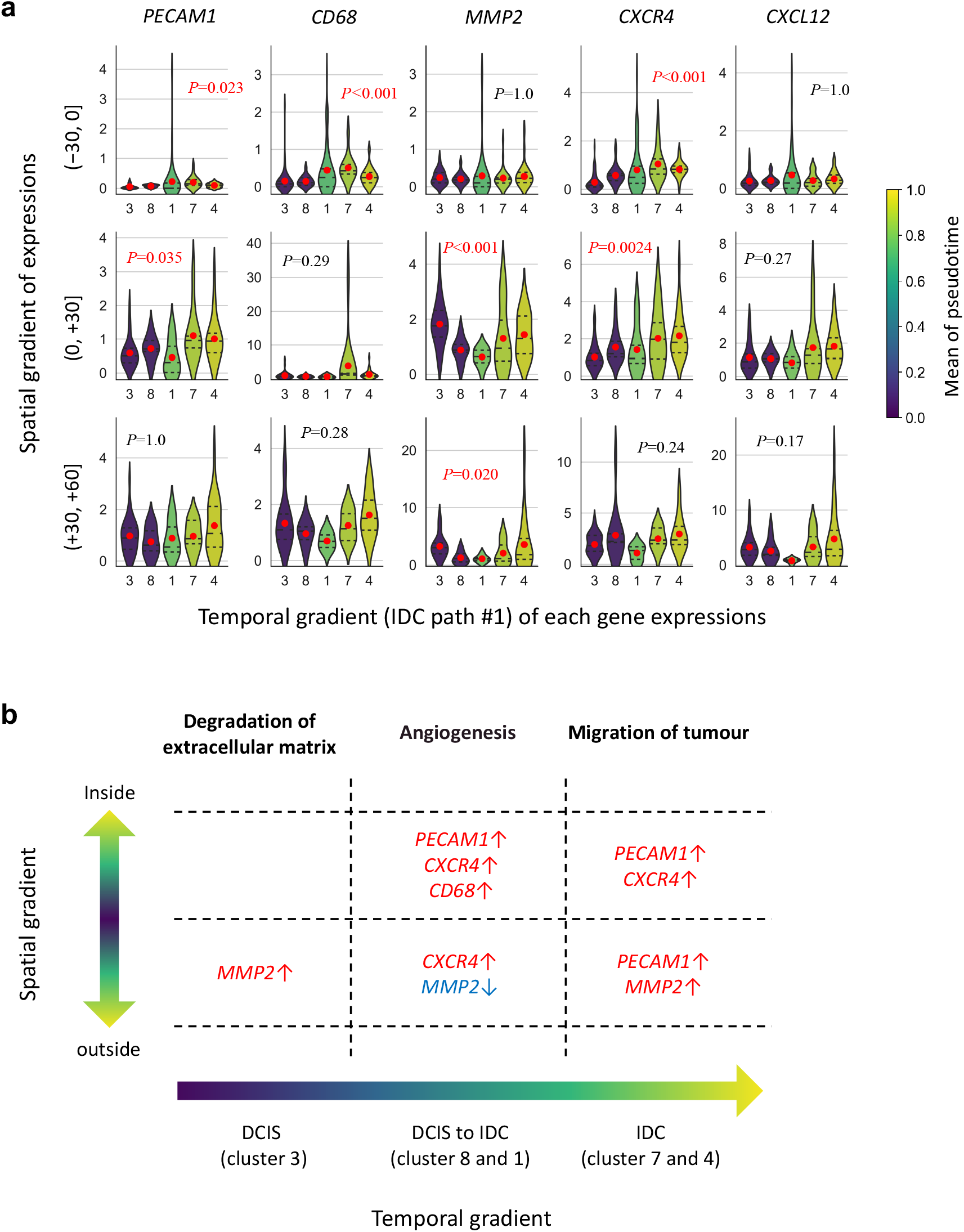
Spatiotemporal trajectory analysis illustrating the progression flow of the tumour microenvironment. **(a)** Violin plots showing the marker expressions of *PECAM1*, *CD68*, *MMP2*, *CXCR4*, and *CXCL12* on the estimated trajectory path (IDC path #1) in the (−30, 0], (0, +30], and (+30, +60] sections. The color scale indicates the mean of pseudotimes in each cluster. The annotated values represent the *P* values of the significance test. The red dots in the figures indicate the mean of the gene expression level. (b) Summary of the dynamics of the gene expressions. The red and blue colours indicate the overrepresentation and underrepresentation of the gene expressions.

Finally, we summarised the temporal sequences of the expression of these genes. In the early stages of cancer progression, the expression of *MMP2* was upregulated in the peri-tumour and outer regions (Fig. 6b). In the tumour progression from non-invasive to invasive cancer, infiltration of endothelial cells (*PECAM1*) and macrophages (*CD68*) was noted into the tumour interior, together with increased chemokine signalling (*CXCR4*). After invasion, the expression of *MMP2* was upregulated in the peritumour and outer regions. Therefore, we concluded that the SKNY showed spatiotemporal sequences of interactions between tumours and other components within the microenvironment.

## Discussion

In this study, we applied the SKNY algorithm to Xenium data extracted from breast cancer to estimate the cellular and molecular functions and mechanisms in the TME. The TME includes diverse cells, such as cancer-associated fibroblasts, stromal cells, and immune cells involved in cancer progression^27^, and the concept of the TME has also been incorporated into clinical research on breast cancer^28^. For example, immunohistochemical pathological analysis has shown that intratumoural macrophages stained by CD68 correlate with malignancy^29, 30^ and that intertumoral microvessel density assessed by CD31, which reflects angiogenesis, is an important poor prognostic factor^31, 32^. In breast cancer, high expression of *Ki67* and *HER2* is associated with malignancy^33^, whereas destruction of myoepithelial cells is associated with tumour invasion^34^. Consistent with these previous reports on the pathology, the results of *spatial stratification* (Output 4) and *spatiotemporal trajectory* (Output 5) analyses, which showed an overrepresentation of *CD68* and *PECAM*1 (*CD31*) within the spatial domain of the invasive tumour (Fig. 5 and 6), demonstrated the infiltration of macrophages and endothelial cells into malignant cancer. Moreover, *MMP2* was overexpressed in the early and late stages of tumour progression in the stromal area, and *CXCR4* and *CXCL12* were enriched after mid-stage progression inside the tumour (Fig. 6). MMPs contribute to the sprouting of vascular endothelial cells by degrading the vascular basement membrane and extracellular matrix in the early stages of tumour angiogenesis^35^, and CXCR4/CXCL12 signalling pathway mediates cell migration signals and metastasis processes^36^. These results are consistent with the previous findings, suggesting that our algorithm can accurately estimate compatible biological mechanisms in the TME.

The *trajectory estimation* (Output 3) analysis was used to construct the tumour progression trajectory of the spatial domains (Fig. 4). The interaction of various cells in the TME is considered crucial in cancer progression^27^; therefore, the progression trajectory should be determined by integrating all cells in the TME rather than focusing solely on cancer cells. Our results showed that during the transition from DCIS to IDC, an over-representation of vascular endothelial cells expressing *PECAM1* and *VWF*, as well as an increase in the *CXCL12* and *CXCR4* chemokine-chemokine receptor pair was noted. These results are consistent with the known mechanisms by which cancer cells acquire invasive potential through endothelial cells^37^ and the associated induction of cell migration signals from chemokines^36^. Most importantly, gene expression from non-cancer cells was the ‘missing link’ between DCIS and IDC in the trajectory, suggesting the utility of this approach to integrate all cells within the spatial domain. Furthermore, our data estimated the trajectory from the root to *PGR*-positive DCIS without progression to IDC. Reduced *PGR* expression has been suggested as a surrogate marker for *GATA3* mutations, one of the genetic factors involved in the progression of DCIS^38,39^. Paradoxically, these previous reports, combined with our results, suggest that the transition to *PGR*-positive DCIS may slow cancer progression. The thin edge from *PGR*-positive DCIS to other clusters in the PAGA graph also supports this hypothesis.

The *detection* algorithm (Output 1) delineated different tumour shapes based on histological features (Fig. 2 and 3). The enrichment of cancer cells and stromal markers within and outside the spatial domains indicates accurate separation of the tumour and stroma. Myoepithelial cells surround the ductal epithelium for structural support^40^, and our results also showed that myoepithelial cell markers, including *ACTA2*, *MYLK*, and *KRT14*, were enriched in the perispatial domain of the tumour, suggesting high-quality detection of tumour contours using our algorithm. This high-quality contour guaranteed subsequent SKNY analyses, including *clustering* (Output 2), *trajectory estimation* (Output 3), and extension into the microenvironment (Outputs 4 and 5).

This study had some limitations. First, we used only one sample for this analysis, and whether the SKNY would work with other samples remains to be determined. Although our preliminary analysis of lung, kidney, colon, and skin samples confirmed their quality (confidential), it is necessary to verify the performance of the SKNY using a large number of samples. Second, in this analysis, the spatial omics data was converted to grids of 10 × 10 μm, and this conversion may make it difficult to detect thin tissues, such as monolayered epithelium. However, setting the grid data to a smaller size should result in insufficient sensitivity of the marker genes on each grid. Therefore, it is necessary to consider the balance between grid size and marker gene sensitivity for each specimen and gene panel.

In conclusion, SKNY can be used in microenvironmental analyses to provide valuable insights into its pathological functions. It should be applicable not only to the TME but also to a wide range of microenvironments, such as tertiary lymphoid structures and myocardial and neuronal microenvironments.

## Methods

### Data acquisition and pre-processing

Breast cancer data from Xenium were downloaded from a public repository (https://www.10xgenomics.com/jp/products/xenium-in-situ/preview-dataset-human-breast). The ‘ReadXenium’ function from stlearn (v0.4.12) was utilised to read the HE images (https://www.dropbox.com/s/th6tqqgbv27o3fk/CS1384_post-CS0_H%26E_S1A_RGB-shlee-crop.png?dl=1) and files containing gene expression and cell coordinates (Xenium_FFPE_Human_Breast_Cancer_Rep1_cell_feature_matrix.h5 and Xenium_FFPE_Human_Breast_Cancer_Rep1_cells.csv.gz). The ‘tl.cci.grid’ function in stlearn was used to simplify the coordinate data into grid data (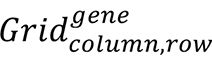 𝑔𝑒𝑛𝑒 = {𝐴𝐵𝐶𝐶11, 𝐴𝐶𝑇𝐴2, 𝐴𝐶𝑇𝐺2, …, 𝑍𝑁𝐹562}, 𝑐𝑜𝑙𝑢𝑚𝑛 = {1,2,3, …, 752}, 𝑟𝑜𝑤 = {1,2,3, …, 547}) at the interval of 10 µm.

### Detection of spatial domain

The pre-spatial domain (𝑆_𝑝𝑟𝑒_) was determined based on the expression of *CDH1* in each grid. The SKNY program can detect prespatial domains based on user selection. For example, a tumour is detected but normal epithelium is not detected upon logical subtraction between a positive marker (e.g. *CDH1*) and a negative marker (e.g. *SFTPB*) expression, which is described as follows:

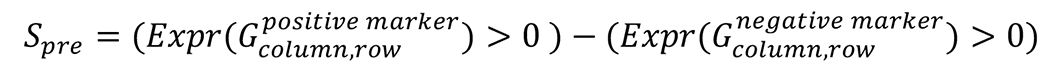

where *Expr* is defined as a function of extracting gene expression counts from the grid. To remove noise from the pre-spatial domain, the "medianBlur" function (kernel size: 3×3) from the Python library opencv (v4.8.1) was applied, resulting in the formation of a spatial domain (*S*) (Supplementary Material 1).

The STARGATE algorithm^24^ was also used to extract spatial domain clusters for comparison with the existing methods. To annotate the extracted spatial domain clusters, the expression levels of epithelial markers (*CDH1*, *EPCAM*) were compared, and cluster 1, 3, and 9, which showed overexpression, was extracted as the spatial domain of the tumour. To assess the concordance between SKNY and STARGATE in the spatial domains, the Jaccard coefficient, which indicates the percentage of agreement between each lattice, was calculated.

### Measurement of distance from the spatial domain surfaces

The edge grids were identified using the’ findContours’ function from opencv in the spatial domain. All adjacent grids were connected by edges and weighted according to the Euclidean distance: 1 for vertical and horizontal edges and 2 for diagonal edges (Supplementary Material 2). The shortest path from the edge grids to the other grids was measured using the multi-source Dijkstra method^41^ to determine the distance from the spatial domain edges.

### Segmentation from a spatial domain to individual spatial domains

The function ‘connectedComponentsWithStats’ from opencv was used to divide the spatial domain (*S*) into individual spatial domains (𝑆_𝑑_, 𝑑 = {1,2,3, …, 426}) with an area larger than 1000 μm^2^. The gene expression within each spatial domain was averaged.

### Spatial stratification by spatial domains

Using the measured distances, a stratification was performed with a half-open interval of 30 µm to obtain the edge grid of the spatial domain (𝑃_𝑥<𝜇≤𝑥+30_, 𝑥 = {0,30}). The rectangle that enclosed each 𝑆_𝑑_ was then extracted, and the contour was enlarged by x μm to produce a rectangle including each spatial domain (𝑅_𝑑,𝑥<𝜇≤𝑥+30_, 𝑥 = {0,30}, 𝑑 = {1,2,3, …, 426}). The peri-spatial domain exclusive to each spatial domain 𝑃𝑆_𝑑,𝑥<𝜇≤𝑥+30_ was calculated as follows:

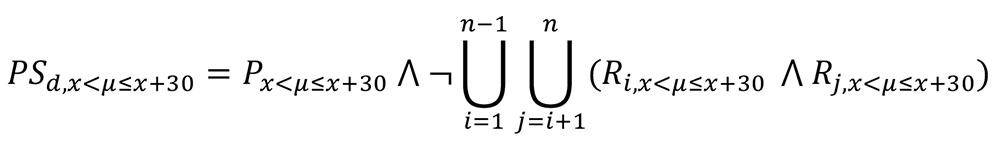

where ⋀ represents the product sum, and ⋃ represents the union set. The gene expression of each 𝑃𝑆_𝑑,𝑥<𝜇≤𝑥+30_ was defined as the average gene expression of the grids within it.

### Diversity analysis in the spatial domain

To compare the alpha diversity of gene expression between the segmented spatial domains and previously annotated cancer cells^26^, the ‘diversity.alpha.chao1’ in the Python library scikit-bio was used to calculate Chao1^42^.

### Clustering of the spatial domains

The ‘pp.log1p’ function from scanpy (v 1.9.8) was used to log-transform gene expression in each spatial domain (𝑆_𝑑_). Then, the ‘pp.pca’ function was used for dimension reduction through principal component analysis. Fifty principal components were extracted in the order of highest eigenvector. The ‘pp.neighbours’ and ‘tl.leiden’ functions from the scanpy were adapted to form spatial domain clusters for Leiden clustering. The function ‘tl.umap’ was used to place leiden embeddings on the UMAP two-dimensional space.

### Trajectory estimation of the spatial domains

For trajectory inference by the PAGA algorithm (ref), the "tl.paga" function of spanpy was used to construct the neighbourhood graph of the spatial domain cluster, followed by the estimation of the pseudotime by adapting the "tl.dpt" function.

### Statistical analysis

Pearson’s product-moment correlation coefficient was used to analyse the correlation between the pseudotime and gene expression. Welch’s t-test was used to compare alpha diversity between the two groups. The Kruskal-Wallis test was used to compare gene expression between multiple groups.

## Visualization

For the visualisation of the Xenium data in space, Xenium Explororor (v1.3) or the "pl.gene_plot" function in stlearn was used.

## Code availability

The code used in this study has been deposited in the documentation of the SKNY library (https://skny.readthedocs.io/en/latest/notebooks/single-TME_analysis.html).

## Acknowledgements

JST SPRING JPMJSP2108.

## Author Contributions

Conception: SA.S., R.Y. and S-I.K.

Construction of algorithm: SA.S.

Data analysis: SA.S.

Validation analysis: R.N.

Implementation of Python library: SA.S.

Pathological consultation: S.N. and S-I.K.

Algorithm consultation: R.Y., Y.S., and A.S.

Discussion per week: SA. S., R.Y., S-I.K., K.T., R.N., and S.C.

Writing original manuscript: SA.S.

Manuscript revision: R.Y., S-I. K., K.T., S.N., Y.S., A.S., R.N., and S.C.

Research supervision (corresponding): S-I.K. and R.Y.

## Competing Interests

The authors report no competing interests.

---

**Supplementary Fig. 1.**
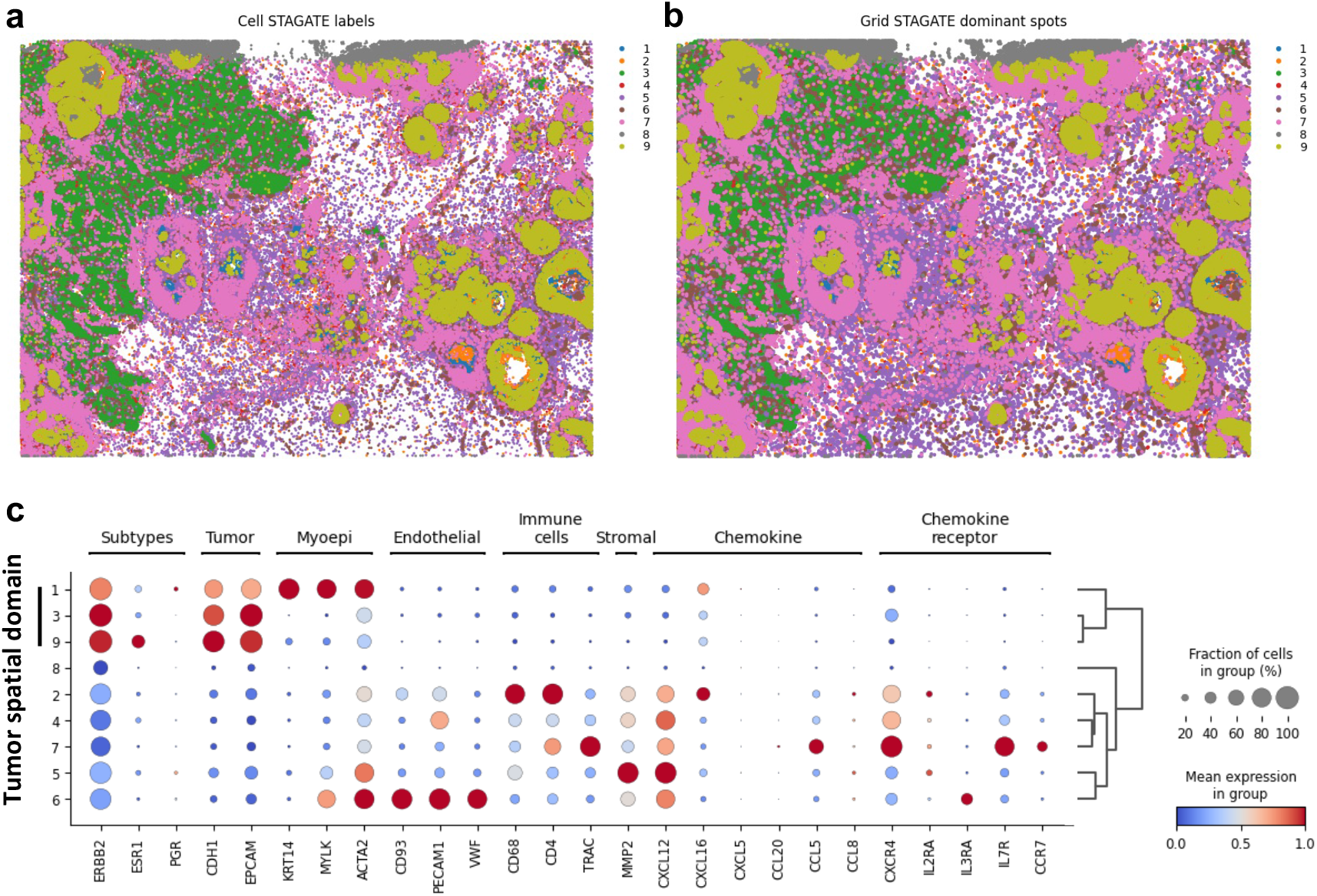
Annotation of the spatial domain using the STARGATE algorithm. Spatial distribution of each cluster by STARGATE algorithm at **(a)** single-cell level and **(b)** grid level. **(c)** Dotplot showing markers of cell types and expression patterns of genes associated with tumour subtypes. Clusters 1, 3, and 9 correspond to the tumour spatial domain.

**Supplementary Fig. 2.**
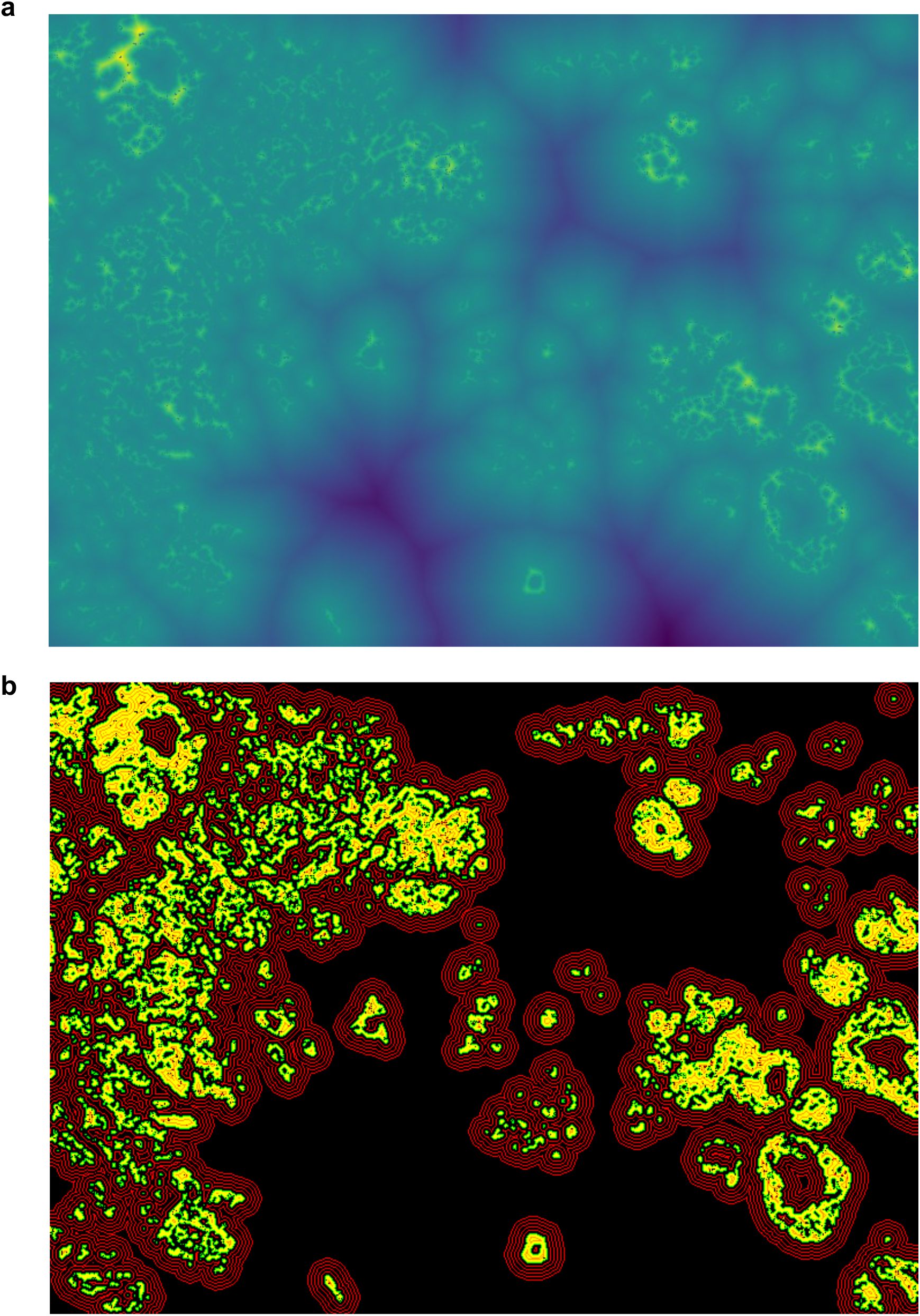
Measurement of distance from the surface of spatial domains. **(a)** Heatmap indicating distance from surfaces of spatial domains. **(b)** The red contour lines indicate distance from the surface of spatial domains at the interval of 30 μm.

**Supplementary Fig. 3.**
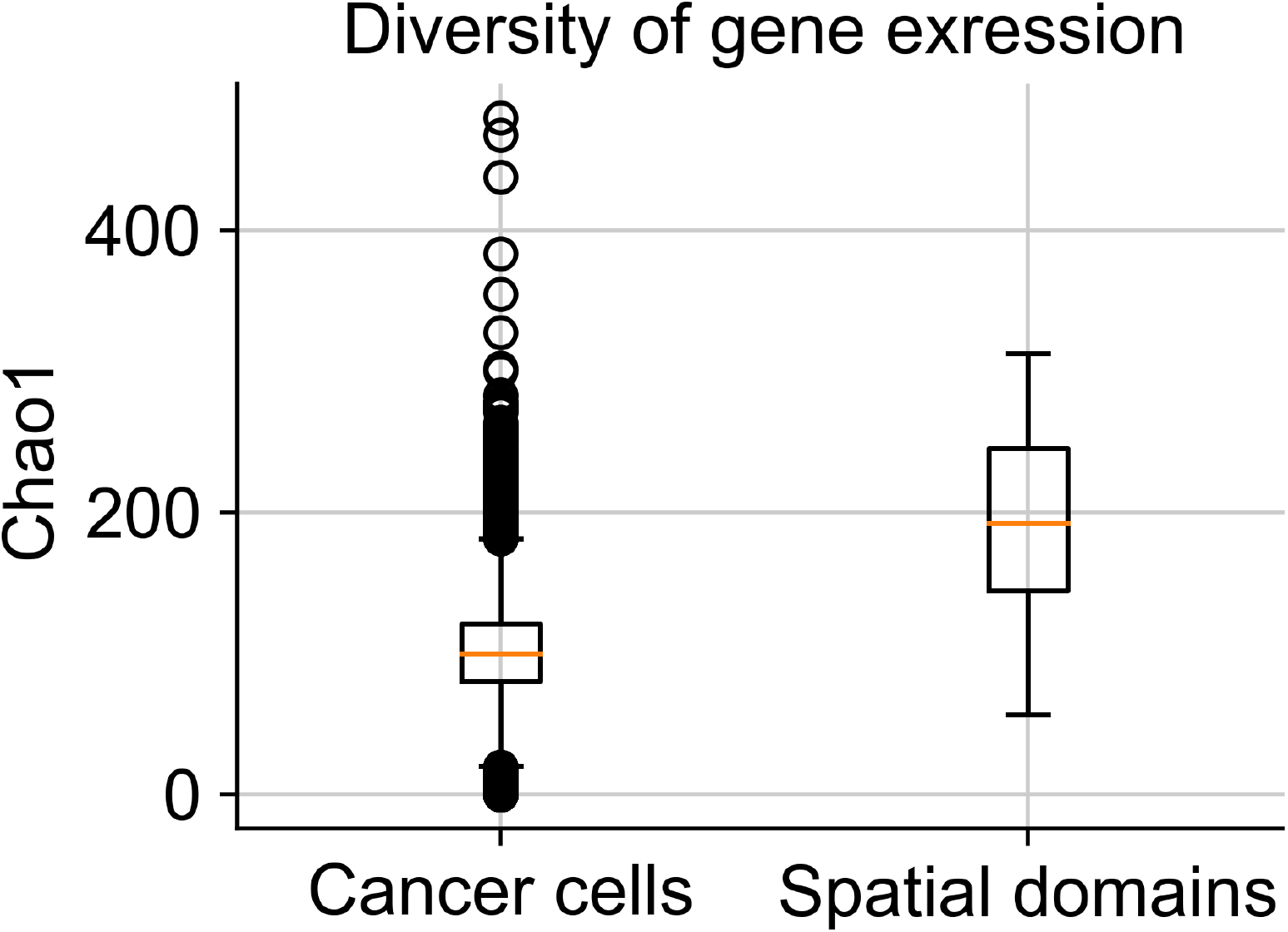
Comparison of alpha-diversity index based on gene expression. Box plot of alpha-diversity index between cancer cells and spatial domains.

**Supplementary Material 1.**
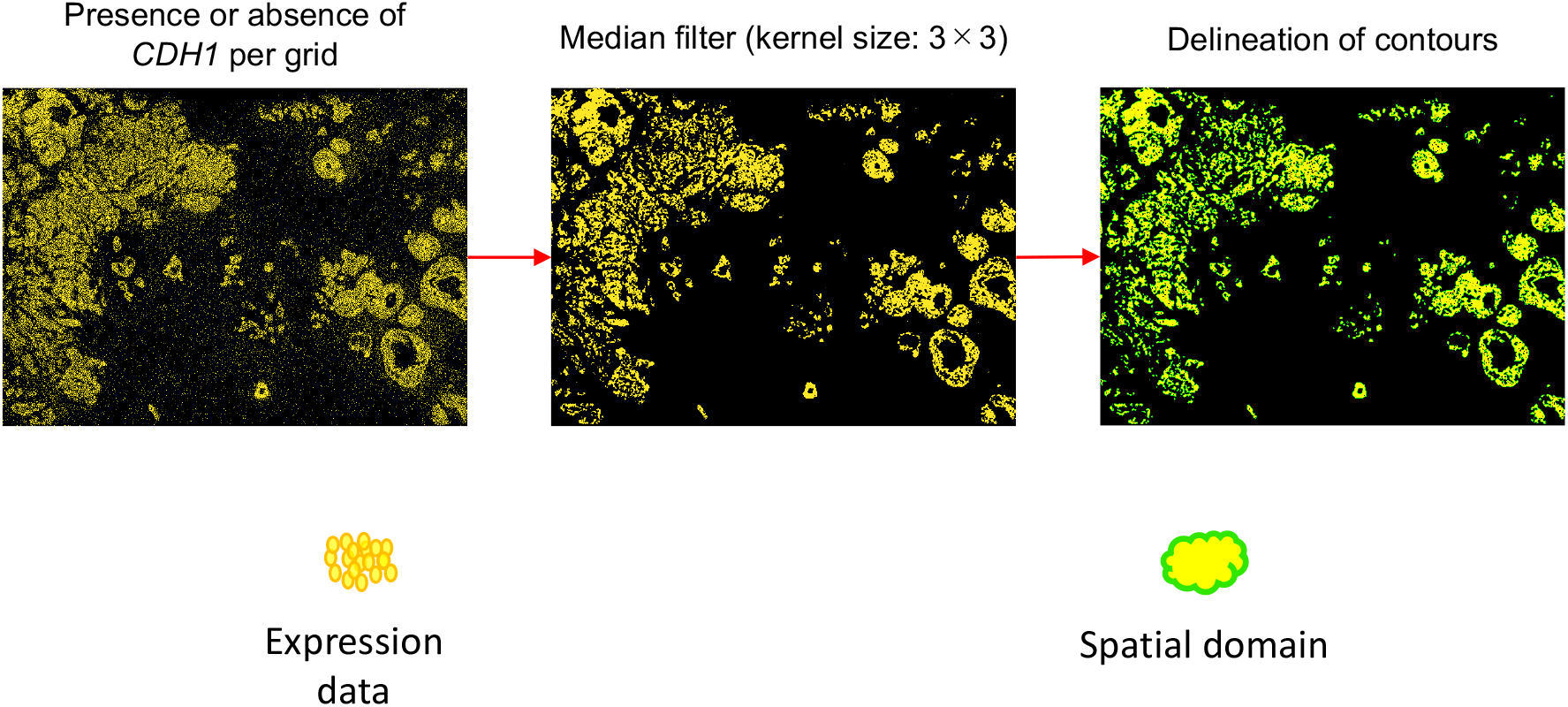
Generation of spatial domain by image processing of gene expression data.

**Supplementary Material 2.**
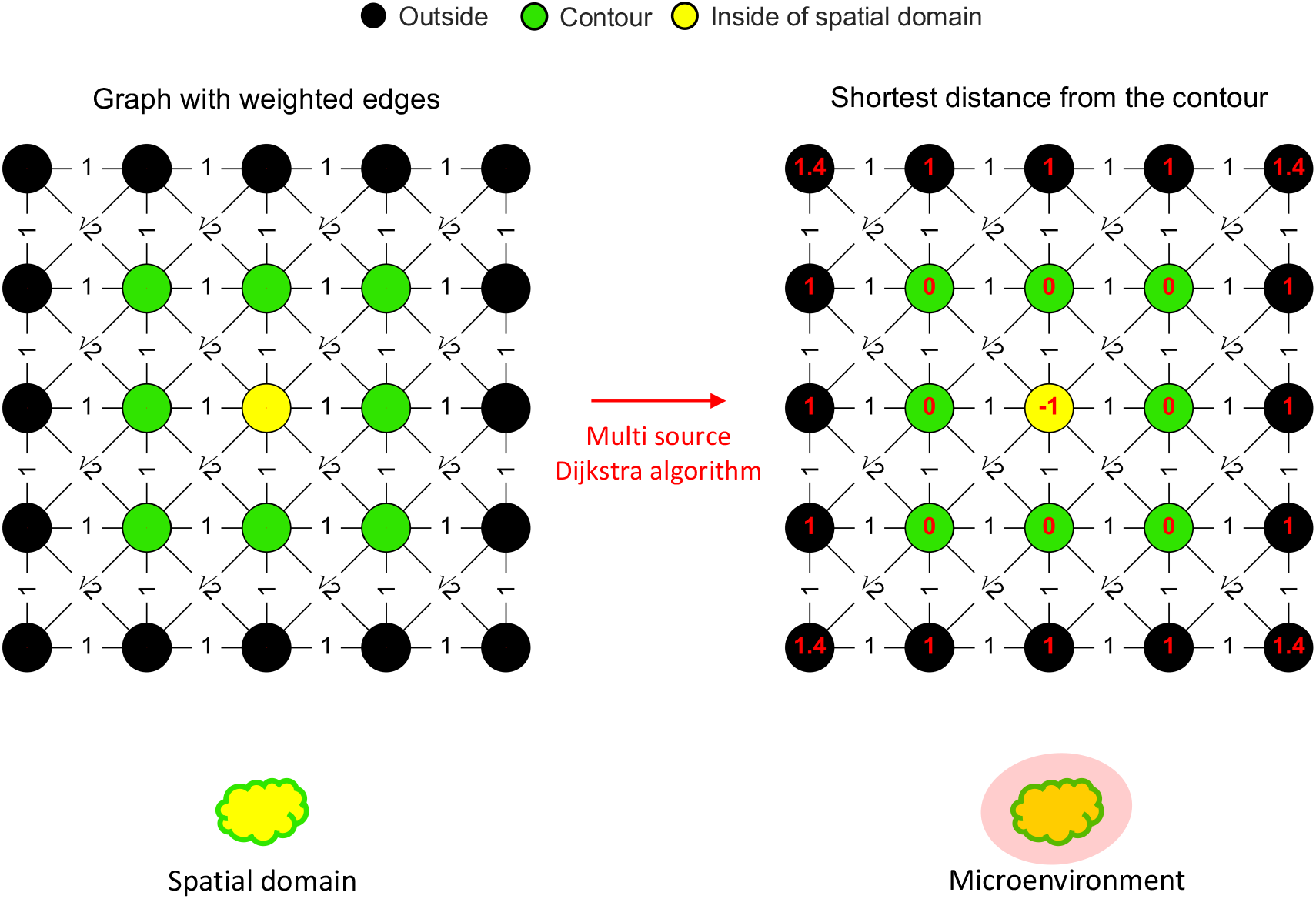
Measurement of shortest distance from contour of spatial domains.

## Notes

### Competing Interest Statement

The authors have declared no competing interest.

